# Limitations of pupil diameter as a proxy for modes of locus coeruleus activity

**DOI:** 10.64898/2026.07.14.738598

**Authors:** Lowell W. Thompson, Siddhartha Joshi, Joshua I. Gold

## Abstract

The locus coeruleus-norepinephrine (LC-NE) system plays multiple roles in higher brain function that are thought to depend on its mode of activation, which reflects relationships between baseline and evoked activation levels. These relationships are evident in both single-unit LC activity and proposed physiological proxies of LC-NE activity, such as pupil size. Here we used measurements in awake monkeys to show that the baseline-evoked relationships evident within these two different measures are unreliably coupled between them: baseline-evoked relationships of the pupil are not predictive of those in the LC, and vice versa. These results imply that pupil modulations, which can reflect LC-NE activity, should be used with caution to make inferences about “phasic” (moderate baseline, high evoked) and “tonic” (high baseline, low evoked) LC-NE activity modes that are thought to support different forms of information processing in the brain.

## Introduction

Cognitive performance follows an inverted-U relationship with arousal (Yerkes and Dodson 1908; Hebb 1955). This relationship is thought to be driven, at least in part, by different patterns of activation of the locus coeruleus-norepinephrine (LC-NE) neuromodulatory system (Aston-Jones and Cohen 2005; Cools and Arnsten 2022). According to this hypothesis, baseline LC activity tracks overall arousal level and shapes the strength of task-evoked LC responses (including “phasic” and “tonic” modes of activation), which in turn influence attention, memory, and other higher-order functions (**Figure 1A**). Low arousal (drowsiness) corresponds to low baseline activity, weak evoked responses, and relatively poor performance. Moderate arousal (optimal engagement) corresponds to moderate baseline activity, strong evoked responses, and relatively high performance (phasic mode). High arousal (distractibility) corresponds to high baseline activity, weak evoked responses, and relatively poor performance (tonic mode). Thus, the difference between phasic and tonic mode involves an inverse relationship between baseline (lower for phasic, higher for tonic mode) and evoked (higher for phasic, lower for tonic mode) LC activity.

**Figure 1:**
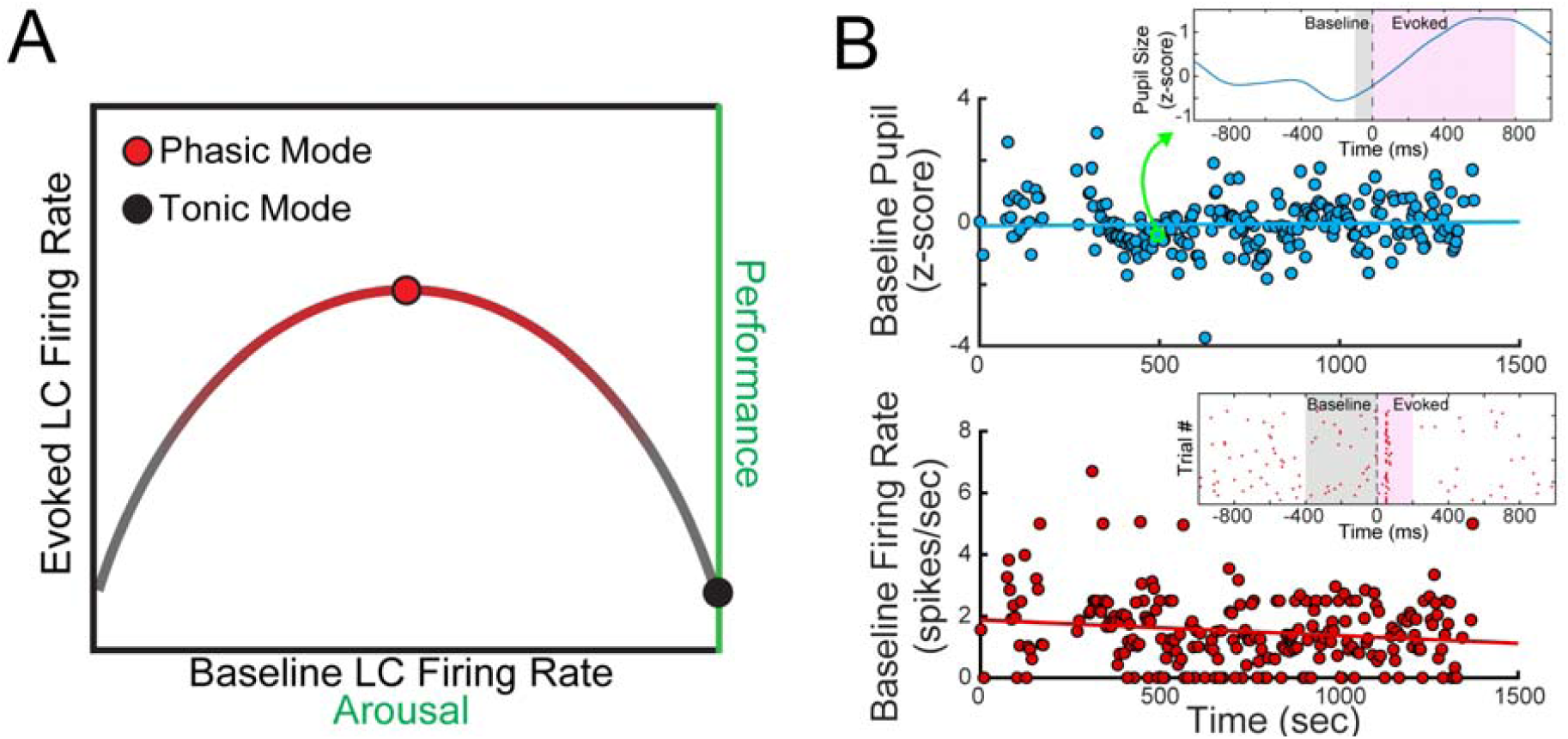
Relationships between arousal, LC activity, and behavior. A) The hypothesized Yerkes-Dodson (inverted-U) relationship between arousal (abscissa, which can be assessed in terms of LC firing rates or non-luminance-mediated changes in pupil diameter) and task performance (right ordinate), with peak performance at moderate levels of arousal. This relationship is thought to reflect specific patterns of LC activation (left ordinate), including task-evoked responses that peak at moderate levels of LC baseline (“phasic mode”) and decline as baseline increases (“tonic mode”). B) Baseline measurements of pupil diameter (top, blue) and the activity of an isolated LC neuron (bottom, red) from a single session. Both were estimated using residuals after accounting for slow (linear) drifts across a recording session (solid lines are linear regression fits). The top inset shows an example pupil response for one trial (green circle) aligned to beep onset. The bottom inset shows an example peristimulus time histogram aligned to beep onset. Shaded regions correspond to the time windows used to quantify baseline (gray) and evoked (magenta) responses.

Experimental support for this hypothesis has been largely indirect. Because the LC is so small and deep in the brainstem, its activity is difficult to measure directly. Thus, studies often rely on proxies. One commonly used proxy is non-luminance-mediated modulations of pupil size, whose relationship to LC is supported by several findings. The LC has direct and indirect projections to circuits that control pupil size (Joshi and Gold 2020). Electrical and optogenetic microstimulation of the LC can evoke pupil responses (Joshi et al. 2016; Y. Liu et al. 2017; Breton-Provencher and Sur 2019; Ghosh and Maunsell 2024; Grimm et al. 2024; Weiss et al. 2026). Pupil changes also track other measures thought to reflect LC activation, including changes in functional connectivity, LC BOLD responses, and certain EEG potentials (Eldar et al. 2013; Murphy et al. 2011, 2014). Moreover, under certain conditions, baseline and evoked pupil size covary with baseline and evoked LC activity, respectively (Joshi et al. 2016).

Despite these findings, many questions remain about the reliability with which pupil measures can be used to infer LC activity. Baseline pupil size is particularly difficult to interpret because it depends on camera position, ambient lighting, and other factors that can vary considerably over hours of recording and across sessions (Mathôt et al. 2018). In contrast, transient responses, which are measured relative to a local baseline, are less vulnerable to these confounds and thus used more often. Some studies have gone further, using evoked pupil responses to infer baseline LC activity, which is central to theories of LC function but difficult to measure (Koss 1986; Gilzenrat et al. 2010; Eldar et al. 2013; Unsworth and Robison 2018). This cross-modal, cross-epoch inference relies on an assumed parallelism between the two measures: in both cases, evoked responses tend to diminish as baseline increases (Aston-Jones and Cohen 2005; Gilzenrat et al. 2010). Thus, a small evoked pupil response is taken to indicate high baseline LC activity (tonic mode), and a large evoked pupil response is taken to indicate moderate baseline activity (phasic mode).

However, this kind of cross-modal, cross-epoch inference about LC firing modes is likely to be complicated by several factors. Pupil size is modulated by multiple neural systems, not just the LC (Joshi and Gold 2020), and the LC’s functional contributions may be modular and cell-specific (Uematsu et al. 2017; Breton-Provencher and Sur 2019; Su et al. 2026). Moreover, the inverse relationship between baseline and evoked pupil size likely reflects not only neural drive but also the pupil’s limited physical dynamic range, which causes response saturation at large baseline diameters (Chen and Kardon 2013). Thus, more direct measurements are needed to better understand if and how pupil-size modulations can be interpreted in terms of phasic and tonic modes of LC activation.

Here we report such cross-modal measurements and show that they should be interpreted with caution. We first replicated previously reported within-epoch relationships between baseline LC activity and pupil size and between evoked LC activity and pupil size, as well as cross-epoch relationships within a single modality. Despite these well-established features of pupil and LC activation patterns, we found no reliable relationships across both modalities and epochs (i.e., baseline LC vs. evoked pupil or baseline pupil vs. evoked LC). These findings further clarify the limitations inherent to interpreting patterns of pupil modulation in terms of neural activity in the LC and elsewhere in the brain.

## Results

We compared measurements of baseline and evoked LC spiking activity (N = 42 single units from 32 sessions in monkey Oz and 63 units from 51 sessions in monkey Ci; an example session is shown in **Figure 1B**) and pupil diameter from monkeys performing a passive fixation task with occasional startling beeps (a reanalysis of data from Joshi et al., 2016). All training, surgery, and experimental procedures were in accordance with the National Institutes of Health Guide for the Care and Use of Laboratory Animals and were approved by the University of Pennsylvania Institutional Animal Care and Use Committee (protocol # 803740).

We first recapitulated previous findings showing reliable relationships between LC activity and pupil diameter when measured separately for baseline (during fixation prior to an auditory beep stimulus) or evoked (during fixation just after an auditory beep stimulus) response epochs (four example units/sessions and population summaries are shown in **Figure 2**, rows 1 and 2). Specifically, trial-by-trial measurements of baseline LC activity were positively correlated with simultaneously recorded measurements of baseline pupil diameter over the course of a session (using residuals, after correcting for linear drifts in both over the session to minimize spurious correlations, as illustrated in **Figure 1B**; Wilcoxon rank-sum test for *H_0_*: median partial Spearman correlation coefficient across sessions *ρ* = 0, *p* = 8.6×10^-4^ for monkey Oz, *p* = 9.85×10^-6^ for monkey Ci). Likewise, baseline-subtracted evoked LC responses were positively correlated with baseline-subtracted evoked pupil responses (*p* = 0.0135 for monkey Oz, *p* = 0.0018 for monkey Ci). These results are consistent with those reported previously and illustrate the coupling between LC activity and pupil size when measured simultaneously.

**Figure 2:**
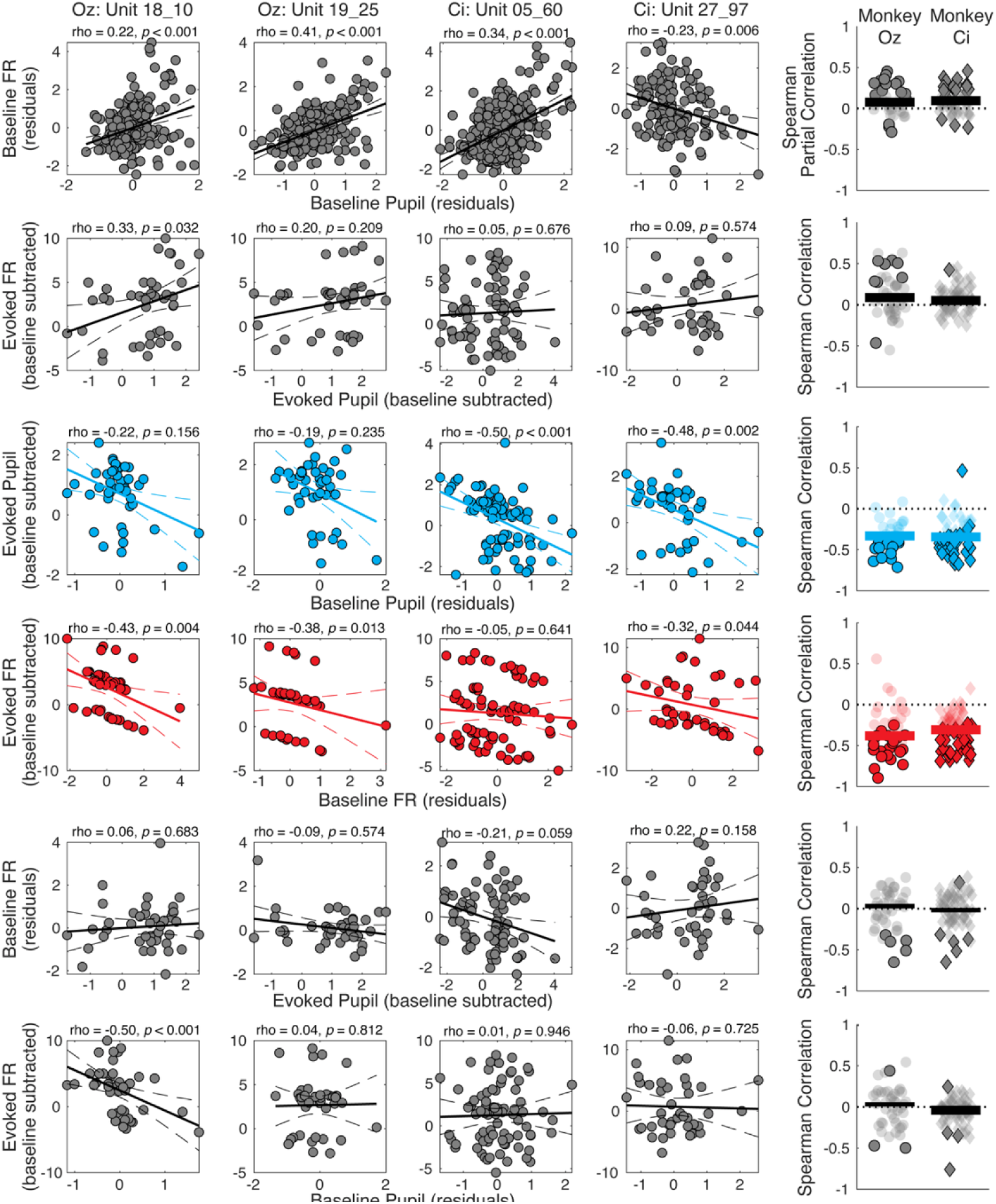
Comparisons of baseline and evoked relationships between LC single-unit activity and pupil diameter. The first four columns show four example units/sessions (in each panel, solid and dashed lines are linear regression fits and 95% confidence intervals). The last column summarizes results across all units/sessions from the two monkeys (darker points indicate units with *H_0_*: partial correlation=0, *p* < 0.05; horizontal bars indicate medians across units, thick bars indicate *H_0_*: median=0, *p* < 0.05). First row: Baseline LC firing rate (FR) versus baseline pupil. Second row: Evoked LC versus evoked pupil. Third row: Evoked pupil versus baseline pupil. Fourth row: Evoked LC versus baseline LC. Fifth row: Baseline LC versus evoked pupil. Sixth row: Evoked LC versus baseline pupil.

During moderate to high levels of arousal, evoked responses are expected to decrease as baseline increases for each respective modality. Accordingly, in our dataset baseline-subtracted evoked pupil responses were negatively correlated with baseline pupil diameter (using residuals; *p* = 1.16×10^-6^ for monkey Oz, *p =* 5.02×10^-9^ for monkey Ci; **Figure 2**, row 3). Similarly, baseline-subtracted evoked LC responses were negatively correlated with baseline LC activity (using residuals; *p* = 5.88×10^-7^ for monkey Oz, *p =* 1.04×10^-11^ for monkey Ci; **Figure 2**, row 4).

Having recapitulated previous findings, we sought to determine whether similar cross-modal (LC and pupil) inferences can be made across different epochs (from baseline to task-evoked and vice versa). We first tested the hypothesis that evoked pupil responses could be used as an inverse approximation of baseline LC activity and corresponding phasic/tonic mode. As shown in **Figure 2** (row 5), evoked pupil responses were not related to baseline LC activity in a reliable manner across sessions (median *ρ* = 0.03, Wilcoxon rank-sum test for *H_0_*: median Spearman correlation coefficient = 0, *p* = 0.894 for monkey Oz; median *ρ* = -0.018, *p* = 0.904 for monkey Ci). Results were similar for the isolated subset of sessions with significant baseline LC-pupil relationships (median *ρ* = 0.005, *p* = 0.642 for monkey Oz; median *ρ* = 0.013, *p* = 0.987 for monkey Ci).

We also tested the complementary hypothesis that baseline pupil diameter could be used as an inverse approximation of evoked LC responses. As shown in **Figure 2** (bottom row), this relationship was relatively small in magnitude for both monkeys, reaching statistical significance for one monkey (Ci), when considering either all sessions (median *ρ* = -0.038, *p* = 0.008) or just those with significant baseline relationships (median *ρ* = -0.069, *p* = 4.98×10^-3^), but in neither case for the other monkey (Oz; full sample: median *ρ* = 0.033, *p* = 0.915; subset: median *ρ* = -0.033, *p* = 0.379). These small, inconsistent across-session trends corresponded to similarly modest within-session effects, with just 4 sessions exhibiting significant relationships for monkey Ci and 3 for monkey Oz. Thus, despite consistent relationships between LC activity and pupil size within each epoch, there were no reliable relationships between the two when comparing across epochs (i.e., baseline versus evoked).

To further examine the reliability and generalizability of these results, we used linear regression with leave-one-out cross-validation for each of the tested models (**Table 1**). The strongest generalizable relationships were for comparisons across epochs for each modality considered separately (i.e., baseline versus evoked pupil and baseline versus evoked LC) or across modalities for each epoch considered separately (i.e., baseline comparisons of pupil versus LC, and substantially less so for evoked pupil versus LC). In contrast, cross-epoch and cross-modal predictions were poor and rarely produced reliable out-of-sample predictions. These results mirror our correlational findings and illustrate that, even when consistent monotonic relationships were identified, they were highly variable and often failed to predict held-out data.

**Table 1.** Cross-Validated Generalization of LC-Pupil Relationships.

| Analysis | CV $R^2$ median | | CV $R^2$ IQR | | CV $R^2$ p-value | | Perm. $p < 0.05$ | |
| --- | --- | --- | --- | --- | --- | --- | --- | --- |
|  | Oz | Ci | Oz | Ci | Oz | Ci | Oz | Ci |
| B. Pupil vs. Ev. Pupil | 0.032 | 0.020 | [-0.138, 0.178] | [-0.123, 0.124] | 0.765 | 0.913 | 16/32 | 26/52 |
| B. LC vs. Ev. LC | -0.047 | -0.048 | [-0.166, 0.059] | [-0.150, 0.006] | 0.105 | <0.001 | 14/42 | 16/63 |
| B. Pupil vs. B. LC | -0.016 | -0.006 | [-0.033, 0.017] | [-0.022, 0.030] | 0.326 | 0.880 | 15/42 | 27/63 |
| Ev. Pupil vs. Ev. LC | -0.134 | -0.086 | [-0.173, -0.047] | [-0.160, -0.049] | <0.001 | <0.001 | 8/42 | 4/63 |
| Ev. Pupil vs. B. LC | -0.106 | -0.081 | [-0.165, -0.067] | [-0.146, -0.045] | <0.001 | <0.001 | 5/42 | 5/63 |
| B. Pupil vs. Ev. LC | -0.131 | -0.100 | [-0.173, -0.069] | [-0.167, -0.054] | <0.001 | <0.001 | 3/42 | 4/63 |

Leave-one-out cross-validation (CV) summary results for each linear regression model (rows). Values are reported across sessions/units. CV $R^2$ compares explained variance on the held-out test set relative to an intercept-only model. Positive values indicate greater variance explained by the complex model compared to the intercept-only model. CV $R^2$ $p$ -values were computed from Wilcoxon rank-sum tests for $H_0$ : median across sessions/units = 0. Permutation tests (perm; $n = 1,000$ ) were conducted for each session/unit to determine the proportion of significant CV $R^2$ values. B = Baseline; Ev = Evoked; IQR = inter-quartile range.

We conducted several additional analyses to address possible sources of these null results. First, the startling auditory beep might have been insufficient to evoke the full dynamic range of LC and pupil responses, potentially limiting the magnitude of the observed correlations. To assess the potential impact of this limitation, we related the correlation coefficients between evoked pupil responses and baseline LC activity (and vice versa) to the range of observed LC firing rates or pupil responses in each session. We found that the relationship between baseline pupil diameter and evoked LC responses was not systematically impacted by the range of evoked LC responses for either monkey (Spearman’s *ρ* = 0.02, *p* = 0.89 for monkey Oz; *ρ* = 0.07, *p* = 0.589 for monkey Ci; **Supplemental Figure 1A**). Similarly, the relationship between baseline LC activity and evoked pupil responses was not systematically impacted by the range of evoked pupil responses for monkey Ci (*ρ* = 0.18, *p =* 0.16; **Supplemental Figure 1A,** green). However, we did observe a modest impact for monkey Oz (*ρ* = -0.33, *p* = 0.03; **Supplemental Figure 1B,** purple), suggesting that evoked pupil diameter may be inversely proportional to baseline LC activity for some conditions that evoke a sufficiently wide range of evoked pupil responses.

Second, our conservative approach to calculating baseline, by regressing out any linear relationship with elapsed time, could eliminate or reduce genuine correlations. However, this effect was minimal in our dataset, and we obtained similar results when using raw baseline measurements as opposed to residuals (see **Supplemental Figure 2**).

Third, it is possible that these relationships are more consistent with a non-linear (e.g., Yerkes-Dodson inverted-U) form that was not well identified by our analysis approaches (e.g., the Spearman correlation coefficient). We therefore fit both linear and quadratic curves for all across-epoch relationships illustrated in **Figure 2** and computed the log-likelihood ratios for the corresponding fits (**Supplemental Figure 3**). The relationship between baseline and evoked pupil responses was better described by a non-linear fit for a minority of sessions (LLR > critical value of 3.841: 7/32 sessions for monkey Oz, 6/51 sessions for monkey Ci). Similarly, LC neuronal responses were rarely better characterized using a non-linear fit (2/42 neurons for monkey Oz, 3/63 for monkey Ci). Accordingly, across-epoch comparisons showed similar trends that favored a linear relationship (baseline LC versus evoked pupil: 4/42 for monkey Oz, 4/63 for monkey Ci; baseline pupil versus evoked LC: 2/42 for monkey Oz, 5/63 for monkey Ci). These results suggest our null findings were not attributable to a potentially non-linear relationship between baseline and evoked responses in our dataset.

**Figure 2 figure supplement 1:**
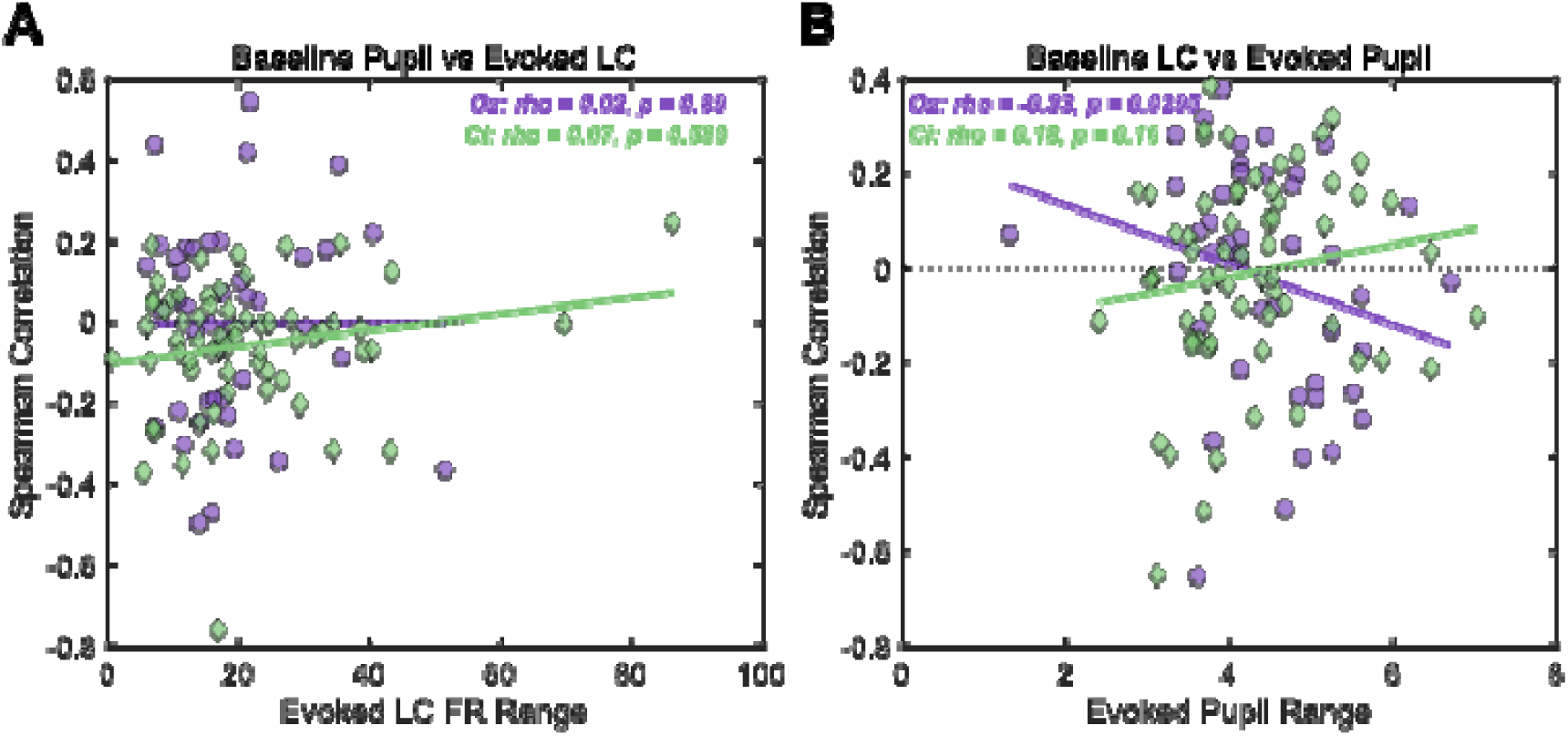
Effects of evoked response range on relationships between LC single-unit activity and pupil diameter. **A)** Effect of evoked LC firing rate range on the relationship between evoked LC and baseline pupil. **B)** Effect of evoked pupil range on the relationship between baseline LC and evoked pupil. Each point is a single unit/session and lines are linear regression fits (purple/circles for Monkey Oz; green/diamonds for Monkey Ci). FR = firing rate.

**Figure 2 figure supplement 2:**
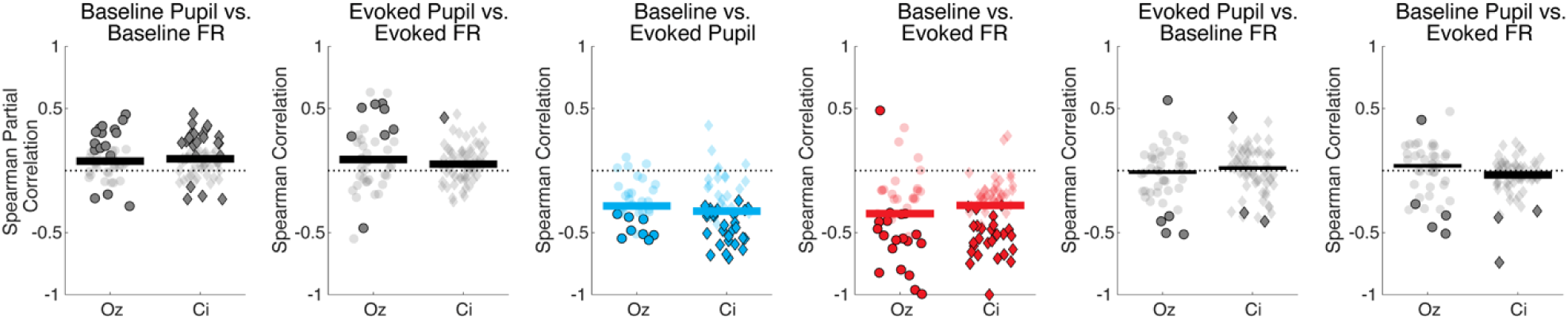
Comparisons of raw baseline and evoked relationships between LC single-unit activity and pupil diameter. Summary of correlation coefficients across all units/sessions from the two monkeys (darker points indicate units with *H_0_*: partial correlation=0, *p* < 0.05; horizontal bars indicate medians across units, thick bars indicate *H_0_*: median=0, *p* < 0.05), plotted as in Figure 2 (last column). Baseline values did not account for potential drifts over time. First column: Baseline LC firing rate (FR) versus baseline pupil. Second column: Evoked LC versus evoked pupil. Third column: Evoked pupil versus baseline pupil. Fourth column: Evoked LC versus baseline LC. Fifth column: Baseline LC versus evoked pupil. Sixth column: Evoked LC versus baseline pupil.

**Figure 2 Figure supplement 3:**
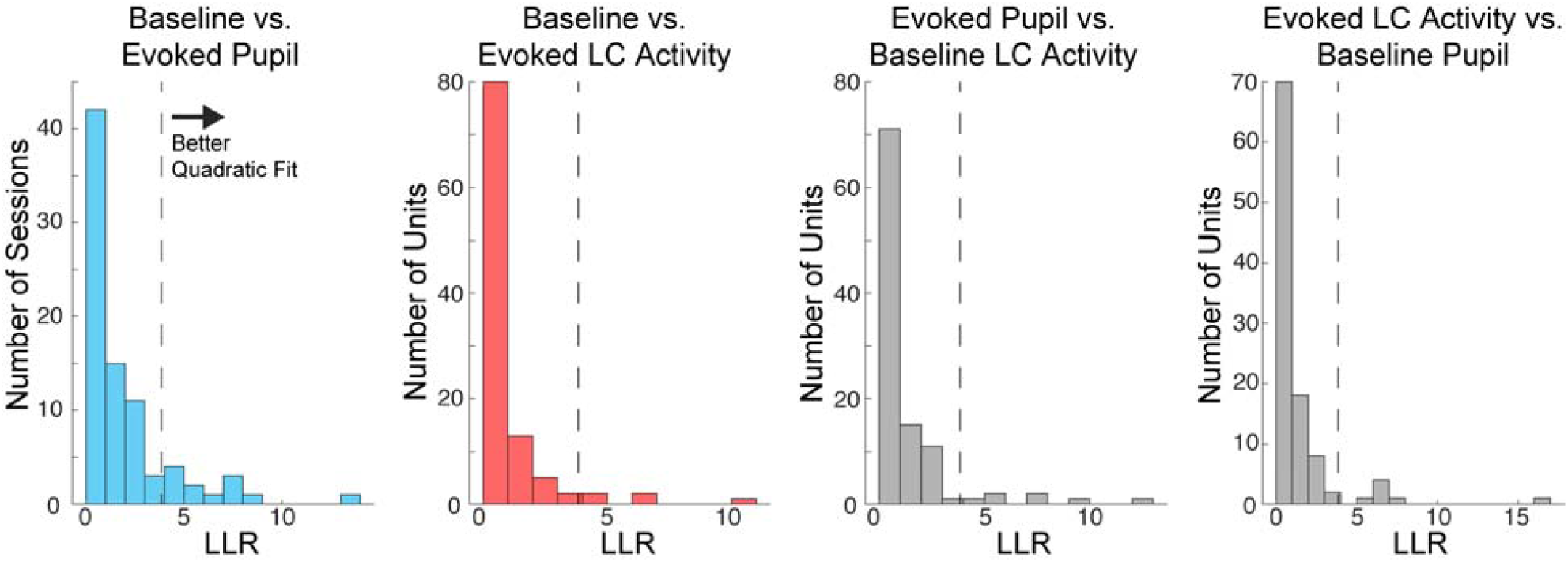
Log-likelihood ratio comparisons of linear and quadratic fits to baseline and evoked relationships. Distribution of log-likelihood ratios comparing the linear and quadratic fits for (from left to right): baseline and evoked pupil responses, baseline and evoked LC activity, evoked pupil responses and baseline LC activity, and evoked LC activity and baseline pupil.

Fourth, the monkeys were exposed to a relatively limited number of trials with startling auditory beeps to evoke LC and pupil responses, which could have underpowered the per-session analyses. To address this possibility, we applied our single-unit analyses (**Figure 2**) to data pooled across all trials, sessions, and units (**Figure 3**). The results are qualitatively similar for the pooled and single-unit analyses, in neither case showing a reliable relationship between baseline pupil and evoked LC responses or baseline LC activity and evoked pupil responses across monkeys. This result also held when LC units were separated based on the direction of their baseline relationship with pupil size (i.e., whether baseline LC activity was reliably associated with larger (**Figure 3**; green) or smaller (**Figure 3**; orange) baseline pupil measurements).

**Figure 3:**
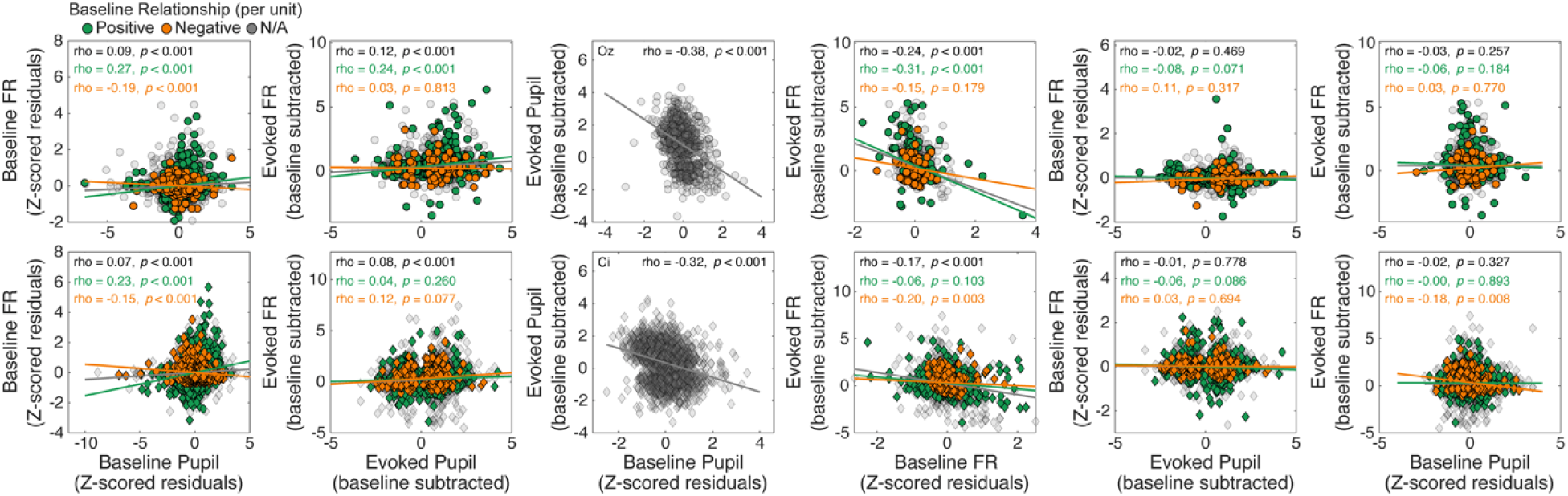
Across-session comparisons of baseline and evoked relationships between neuronal activity and pupillary responses. Measures of LC spiking activity and pupil diameter (z-scored separately for each session/unit) were pooled across trials, sessions, and units for each monkey (rows). First column: Baseline LC firing rate (FR) versus baseline pupil (including non-beep trials). Second column: Evoked LC versus evoked pupil. Third column: Evoked pupil versus baseline pupil. Fourth column: Evoked LC versus baseline LC. Fifth column: Baseline LC versus evoked pupil. Sixth column: Evoked LC versus baseline pupil. Points (data from individual sessions/units) and lines (linear regression fits) are colored based on whether the within-session relationship between baseline pupil diameter and LC activity was significantly positive (green), negative (orange), or was based on all data (grey; see Fig. 2).

## Discussion

Arousal, measured in terms of physiological (e.g., pupil size) or neurophysiological (e.g., the LC-NE system) mechanisms, can interact with higher brain function over multiple timescales. These interactions are often reduced to two primary categories: 1) those that occur over relatively long timescales, which involve fluctuations in arousal that are not tied directly to specific external events but instead represent an internal state that can vary over seconds, minutes, hours, or days (“baseline”) (Cowley et al. 2020); and 2) those that occur over relatively short timescales, which involve abrupt changes in arousal triggered by specific internally or externally generated events that last around a second or less (“evoked”). Both baseline and evoked changes in arousal have been linked to different features of higher brain function (Berridge and Waterhouse 2003; Aston-Jones and Cohen 2005; Cools and Arnsten 2022; Joshi and Gold 2022) and, critically, with each other: evoked changes are often strongest at intermediate baseline values and substantially weaker otherwise, corresponding to the “inverted-U” Yerkes-Dodson relationship. Thus, the role of arousal in higher brain function is often studied and interpreted in terms of coupled variations in baseline and evoked activation patterns.

These studies are dominated by measures of pupil size, which is much easier to measure than LC-NE neural activity (Usher et al. 1999; Gilzenrat et al. 2010; Laeng et al. 2012; Eldar et al. 2013). Because of known relationships between the two, studies that measure pupil size are often interpreted in terms of LC-NE function. Here we reproduced previous findings that support links between the two measurement modalities when considering baseline or evoked activity separately. Specifically, variability in baseline values of LC activity and pupil size measured across trials lasting ∼2–4 s, and variability in evoked changes in LC activity and pupil size in response to a surprising stimulus, each tended to show positive correlations. These relationships were statistically reliable for about half of the recorded neurons, consistent with previous observations in other species (Megemont et al. 2022), aligning with the idea that pupil changes are dynamically coupled with multiple neural systems in a manner that depends on an animals’ cognitive, behavioral, and physiological state (Berridge and Waterhouse 2003; Reimer et al. 2016; Yang et al. 2021; Cazettes et al. 2021; Joshi and Gold 2022; Y. A. Liu et al. 2025; Weiss et al. 2026).

We also examined relationships between baseline and evoked changes within and across each recording modality. Consistent with previous results, we found reliable within-modality relationships: baseline and evoked measurements tended to be negatively correlated with each other when analyzed separately for pupil size or LC activity, such that evoked responses decreased as baseline increased. However, these relationships did not hold across modalities: baseline pupil size did not covary reliably with LC evoked activity, and baseline LC activity did not covary reliably with evoked changes in pupil size.

These findings imply that the two systems are insufficiently coupled to reliably use moment-to-moment changes in pupil size to predict “phasic/tonic” activity modes of individual LC neurons. This limitation likely results from several factors. First, it is possible that the single-unit relationships between the activity of our sample of LC neurons and pupil size differs from the collective output of the area. Moreover, multiple neural systems contribute to pupil changes, and even within the LC there are functional subdivisions with distinct inputs (at least in rodents) that may differ in their relationships to pupil control (Uematsu et al. 2017; Breton-Provencher and Sur 2019; Joshi 2025; Y. A. Liu et al. 2025; Su et al. 2026). The relationship between LC activity and pupil size is also likely impacted by the distinct mechanisms that govern relationships between baseline and evoked activity patterns in the brain and the eye. These mechanisms include spike-generating processes in the LC and physical properties of sphincter muscles in the eye that introduce different saturating non-linearities that tend to reduce evoked responses as baseline increases. Thus, this limitation also likely depends on the ranges of baseline and evoked activity considered. In our study, the monkeys were generally engaged in the task, suggesting that they were operating around the peak of the inverted-U Yerkes-Dodson curve. Targeted experiments aimed at identifying the distinct contributions of different neuromodulatory systems, through both pharmacological and task-specific manipulations, will be crucial in developing a comprehensive understanding of the conditions under which pupil diameter can be used to make reliable inferences about LC-NE function.

## Materials and Methods

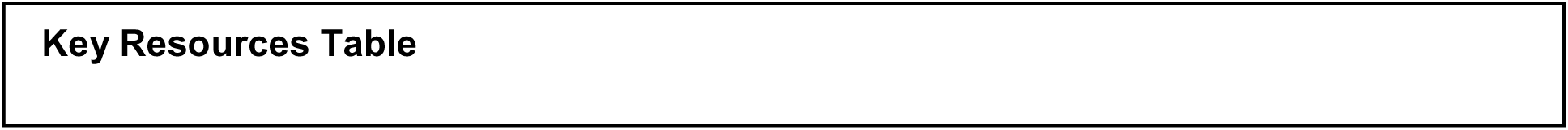
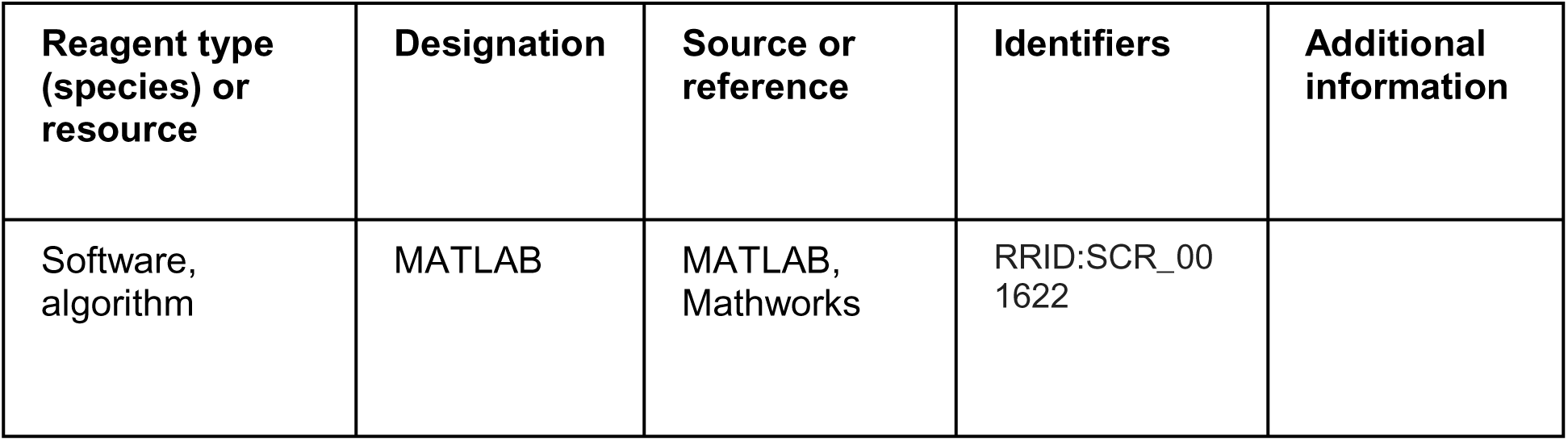

**Key Resources Table**
| Reagent type (species) or resource | Designation | Source or reference | Identifiers | Additional information |
| --- | --- | --- | --- | --- |
| Software, algorithm | MATLAB | MATLAB, Mathworks | RRID:SCR_001622 |  |

### Behavioral Task

The monkeys performed a fixation task that required them to maintain fixation for a variable duration of 1–5 sec. On ∼25% of the trials, selected at random, a loud auditory tone (1 kHz; 0.5 sec) was played after 1–1.5 sec of fixation. Successful fixation until fixation-point offset on beep and no-beep trials was followed by a liquid reward.

### Pupillometry

A video-based eye-tracking system (Eyelink 1000, SR Research) was used to obtain measures of pupil diameter (see Joshi et al. 2016 for details). Briefly, fixation was measured monocularly (1000 Hz sample rate) and only included periods of stable fixation (those without saccadic eye movements) that were maintained for at least 200 ms. Raw pupil measurements were first z-scored for each session. To remove modulations related to fixation onset, the average trace aligned to fixation onset and measured across trials was subtracted from each trial. These standardized traces were then smoothed using a 151-ms wide boxcar filter. Baseline pupil diameter was quantified as the mean smoothed pupil trace throughout fixation for fixation-only trials, or until the onset of an auditory beep (minimum baseline duration of 100 ms) for beep trials. Evoked pupil diameter was quantified as the maximum change in pupil diameter (from mean values in 50 ms-wide sliding windows measured in 10-ms steps) relative to baseline following the onset of an auditory beep until 20 ms before the end of fixation.

### Neuronal Recordings

Details of the neuronal recordings were reported previously (Kalwani et al. 2014; Joshi et al. 2016). Briefly, monkeys were implanted with circular recording cylinders (∼2 cm diameter) that allowed for custom microelectrodes to be lowered approximately dorsal-ventrally while electrical activity was recorded from multiple structures typically encountered leading to the pontine brainstem, including the superior colliculus, inferior colliculus, and the locus coeruleus. Recordings were filtered between 100 Hz and 8 kHz to extract spiking activity and were sorted offline using the Plexon Offline Sorter (Plexon, Inc.). For each single unit, baseline activity was quantified as the firing rate from fixation onset to 20 ms before the end of fixation for fixation-only trials or until the onset of an auditory beep (minimum baseline duration of 100 ms) for beep trials. Evoked activity was quantified as the baseline-subtracted firing rate 0–200 ms following an auditory beep. All statistical analyses were conducted in MATLAB (R2023b; Mathworks).

## Data Availability

All data and analysis code necessary to generate figures is accessible at https://github.com/lwthompson2/Thompson2026_LCPupilAnalysis.git

## Author Details

### Contribution

Conceptualization, Software, Formal analysis, Visualization, Writing – review and editing

### Contribution

Data acquisition, Writing – review and editing

### Contribution

Conceptualization, Software, Formal analysis, Funding acquisition, Visualization, Writing – review and editing

### Ethics

All training, surgery, and experimental procedures were in accordance with the National Institutes of Health Guide for the Care and Use of Laboratory Animals and were approved by the University of Pennsylvania Institutional Animal Care and Use Committee (protocol # 803740).

### Funding Information

National Institute of Mental Health (NIMH): Joshua I. Gold; R01-MH-127566

