## Supplementary figures and images for "Limitations of pupil diameter as a proxy for modes of locus coeruleus activity"

### Supplemental Figure 1

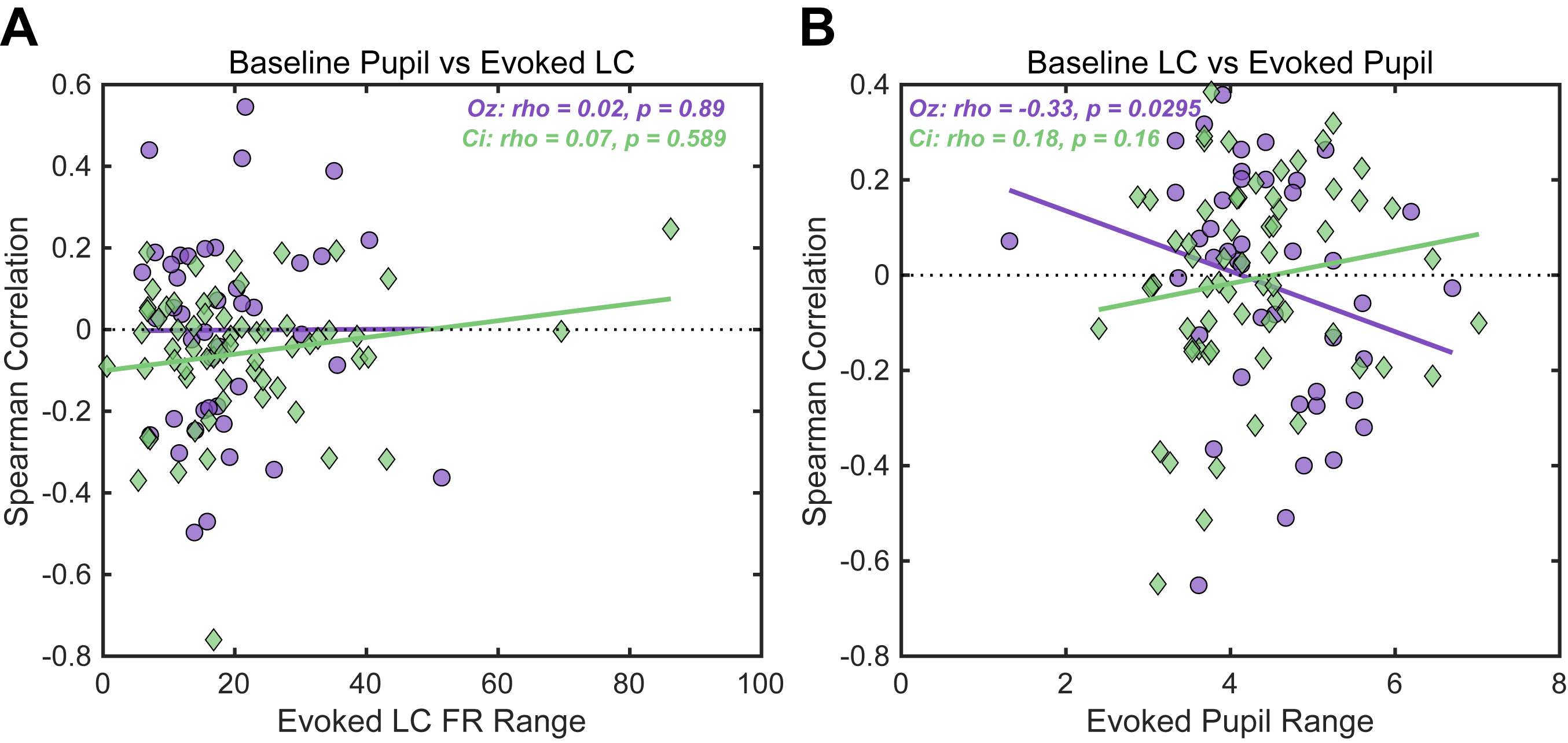

### Supplemental Figure 2

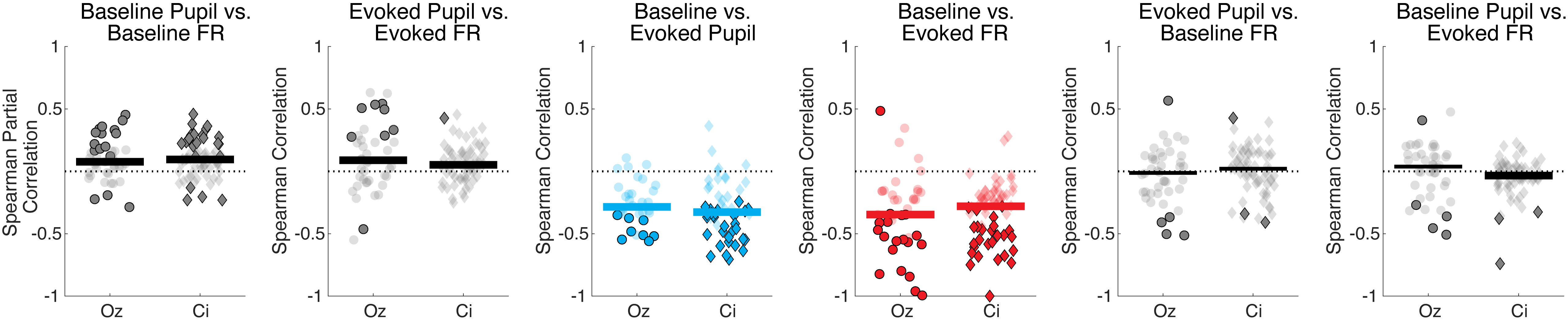

### Supplemental Figure 3

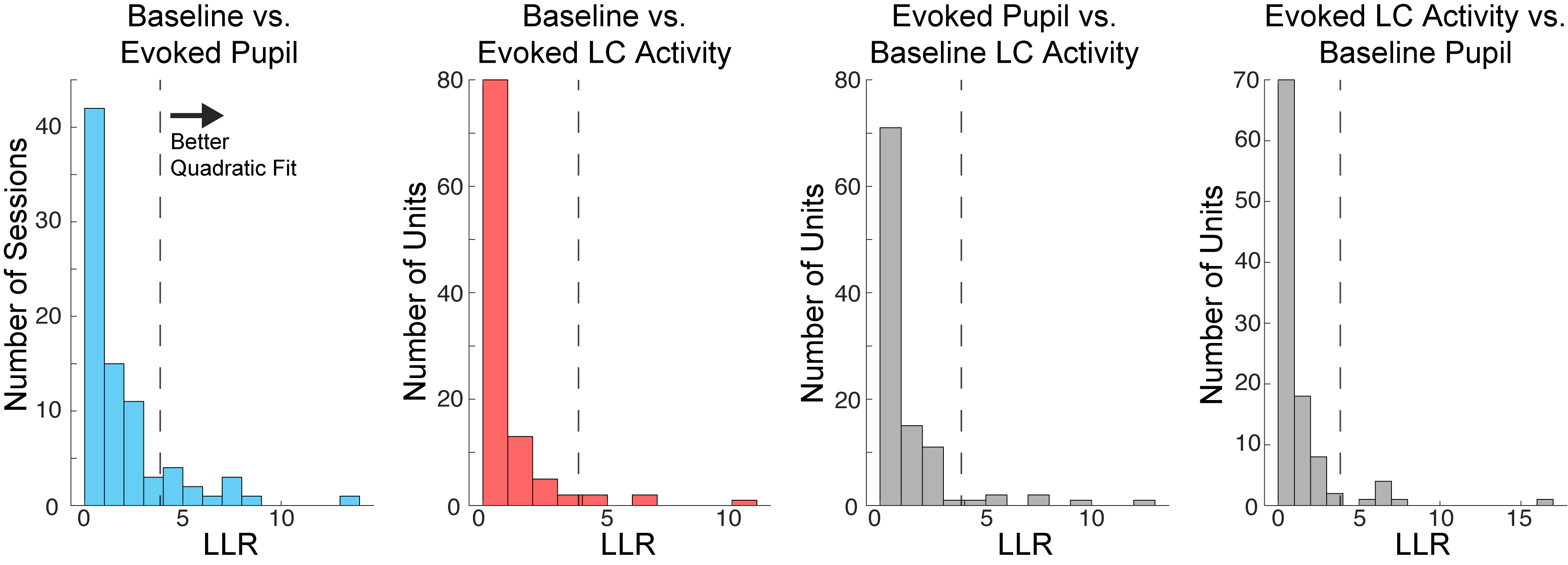
